# Self-propagating wave drives noncanonical antidurotaxis of skull bones in vivo

**DOI:** 10.1101/2023.07.10.547677

**Authors:** Yiteng Dang, Johanna Lattner, Adrian A. Lahola-Chomiak, Diana Alves Afonso, Anna Taubenberger, Elke Ulbricht, Steffen Rulands, Jacqueline M. Tabler

## Abstract

Cell motility is a key feature of tissue morphogenesis, and it is thought to be driven primarily by the active migration of individual cells or collectives. However, this model is unlikely to apply to cells lacking overt cytoskeletal, stable cell-cell or cell-cell adhesions, and molecular polarity, such as mesenchymal cells. Here, by combining a novel imaging pipeline with biophysical modeling, we discover that during skull morphogenesis, a self-generated collagen gradient expands a population of osteoblasts towards a softer matrix. Biomechanical measurements revealed a gradient of stiffness and collagen along which cells move and divide. The moving cells generate an osteogenic front that travels faster than individual tracked cells, indicating that expansion is also driven by cell differentiation. Through biophysical modeling and perturbation experiments, we found that mechanical feedback between stiffness and cell fate drives bone expansion and controls bone size. Our work provides a mechanism for coordinated motion that does not rely upon the cytoskeletal dynamics of cell migration. We term this self-propagating motion down a stiffness gradient, noncanonical antidurotaxis. Identification of alternative mechanisms of cellular motion will help in understanding how directed cellular motility arises in complex environments with inhomogeneous material properties.

## Introduction

Tissue morphogenesis requires the coordinated motion of cells to move into regions where they eventually differentiate into various specialized cell types. Cell migration is typically thought to be driven by external gradients breaking tissue isotropy and introducing a preferred direction along which cells migrate. Such gradients can arise from chemical cues (chemotaxis (*1*)), electrical fields (galvanotaxis (*2*)), or stiffness (durotaxis (*3*)). In all these examples, the motion of cells is driven by the cytoskeletal or membrane dynamics of individual cells, which may be coupled to one another to drive collective cell migration.

Tight mechanical coupling in epithelia allows for the propagation of physical force across collectives to direct tissue deformations such as convergent extension or collective migration (*4, 5*). Cell-cell and cell-substrate adhesions are also essential in the collective migration of the mesenchymal neural crest where contacts with neighbors repolarize cells towards regions of lower density and substrate interactions provide traction (*6, 7*). However, as organs are built later in development, mesenchyme will generate contiguous tissues with limited free space between domains that are not always separated by overt boundaries such as a basement membrane (e.g. cranial mesenchyme). Additionally, the migratory potential of mesenchyme must also be restricted if not inhibited entirely to maintain tissue cohesion (*8,9*). Therefore, cell migration, as currently described, is insufficient to explain motility in all contexts involving polarised growth or coordinated dynamics in cell collectives.

In physical contexts, motility can be generated spontaneously through the physical interactions of objects or molecules. Local temperature increase in gasses, for example, causes pressure gradients that drive the directed motion of molecules towards cooler regions. Such dynamics can be generated in fluids and within cells as well. At the cellular scale, laser-induced heating of cytoplasm can drive spontaneous cytoplasmic streaming that is sufficient to direct cellular polarity in *C. elegans* (*10*). At the tissue scale, spontaneous motion can arise from inhomogeneous proliferation rates resulting in regions of high proliferation exerting pressure on surrounding cells (*11,12*). Similarly, cell motion within intestinal villi is generated by proliferation in the crypt that displaces neighbors due to mechanical coupling through cell-cell adhesions typical of epithelia (*13*). In these systems, spontaneous motion is generated from the inhomogeneity of the surrounding physical structure, rather than the intrinsic dynamic behaviors of each object within the system.

To ask whether mesenchymal motility can be generated spontaneously through physical mechanisms other than cell migration, we turn to the skull. The frontal bones or calvaria are formed by bone rudiments that expand medially from the side of the head towards the skull midline (*14*) between dermal and meningeal layers with which the prospective skull is entirely contiguous. The expansion of these frontal bone rudiments is thought to occur through directed and collective migration of bone-producing osteoblasts towards the top of the head, leaving behind a collagen meshwork that undergoes concomitant mineralization (*15–17*). Such a progressive pattern of mineralization generates material inhomogeneities along the axis of expansion. Here, we show that the feedback between a self-generated stiffness gradient and cell fate drives cell motility and differentiation to coordinate the expansion of mesenchymal bones. We discover a mechanism of mesenchymal cell motion that does not rely on intrinsic cytoskeletal dynamics to drive directed motion, but rather on cells responding to and actively shaping changing material properties as the tissue develops.

## Results

### Inhomogenous structure across the axis of frontal bone expansion influences cell motility and division

We first measured the inhomogeneity of tissue material properties and extracellular matrix across the axis of bone growth during peak stages of skull morphogenesis (Fig. 1A; Fig. S1). Atomic Force Microscopy (AFM) and nanoindentation across three locations along the axis of bone extension (Fig. 1B) revealed a stiffness gradient that was highest in the bone center and lowest in the undifferentiated mesenchyme (Fig. 1C; Fig. S2A). Through Second Harmonic Generation (SHG) imaging, we found that this stiffness gradient coincides with enrichment of fibrillar collagen (Col1a1) within differentiated mesenchyme (Figure 1C-D). These data demonstrate inhomogeneity in tissue material properties and structure during bone morphogenesis.

**Figure 1.**
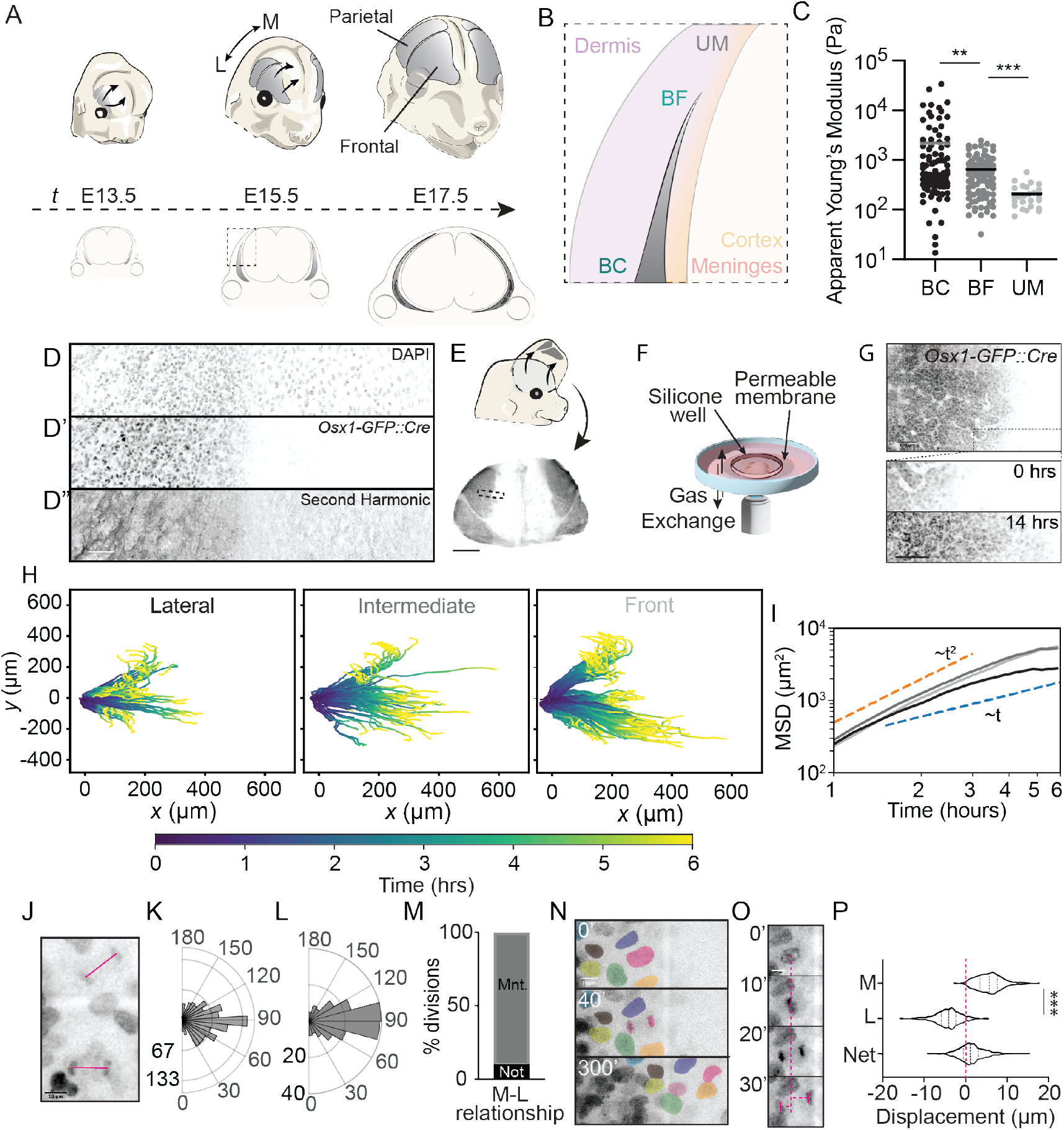
Inhomogeneous material properties and cell behaviors during skull morphogenesis. **(A)** Illustration depicting anisotropic expansion of frontal and parietal bones (grey) toward the top of the head between E13.5, 15.5, and 17.5. **(B)** Diagram depicting a coronal section of the developing skull with the frontal bones labelled in cyan. Dotted box indicates inset showing the locations of the bone center (BC), bone front (BF) and undifferentiated mesenchyme (UM). **(C)** Atomic force microscopy (AFM) measurements of the stiffness (apparent Young’s modulus) at the three locations indicated in **(B)**. Horizontal bars indicate mean. **(D)** Nuclear 540 nm autofluorescence generated by 940 nm excitation of osteogenic front of E14.5 flat-mounted *Osx1:GFP-Cre* skull cap. **(D’)** Osx1::GFP-Cre fluorescence. **(D”)** Second Harmonic Generation signal with 940 nm excitation. **(E)** Schematic showing excision of Sp7-mCherry labelled skull caps. **(F)** Diagram depicting *ex vivo* imaging setup. **(G)** Max projection of *Osx1-GFP::Cre* labelled frontal bone at 0 h at E13.75. Dotted box indicates insets at 0 h and 14 h shown below. Scale bar = 100 mm. **(H)** Graph showing example lateral, intermediate and front cell tracks from live imaging experiments combined (N=4), colour-coded by time. **(I)** Graph showing mean squared displacement for osteoblasts in different mediolateral positions, averaged over N=4 images. Shading indicates SEM. **(J)** Max projection of *Osx1-GFP::Cre* labelled E13.75 live explant showing two late anaphase nuclei with pink lines indicating angle of division. **(K)** Angle of osteoblast division in E13.75 *Osx1-GFP::Cre* labelled live explants, (Kuiper uniformity test *p >* 0.05, *n* = 450, *N* = 8). **(L)** Angle of division orientation in fixed Osx-1::GFP-Cre labelled E13.5 flat-mounted skull caps (Kuiper uniformity test *p >* 0.05, *n* = 202, *N* = 6). **(M)** Graph showing the percentage of osteoblasts that maintain (Maint., grey) or do not maintain (Not maint., black) their mediolateral neighbor relationship. **(N)** Representative stills from *Osx1-GFP::Cre* labelled cultured explants. Osteoblast nuclei have been false colored to demonstrate neighbor relationships, sister nuclei are overlaid with pink. Scale bar = 10 mm. **(O)** Representative time series of an osteoblast division from *Osx1-GFP::Cre* labelled explant showing how nuclear displacement is measured. Scale bar = 10 mm. **(P)** Graph showing significant difference in the displacement of medial and lateral sister nuclei after a division (Mann-Whitney test, *p <* 0.0001).

To test whether cell motion could be affected by these inhomogenous tissue material properties we developed an *ex vivo* skull imaging system. Entire skull caps were explanted from *Osx1-GFP::Cre* mice which harbor an osteoblast-specific GFP::Cre recombinase fusion protein (*18*) (Fig. 1E-G). In this system, cells contiguous with the bone ahead of the front are unlabelled. Although cell membranes are too complex to parse individual cells (Fig. S3A, Supplemental Movie 1), our approach allowed us to distinguish individual nuclei as a proxy for individual osteoblasts for which motion and cell division could be tracked (Fig. S3B,C; Supplemental Movie 2). We tracked individual nuclei at the osteogenic front separating differentiated osteoblasts from undifferentiated mesenchyme, as well as 200 μm and 400 μm towards the bone center (Fig. 1H). While the mean squared displacement (MSD) of all cells initially scales with time as *t*^2^ for all tracked cells indicating ballistic motion, over time the MSD of the cells towards the bone center approaches a linear scaling with time which reflects diffusive motion (Fig. 1I; S4A). In contrast, the cells in the middle and near the front retain their ballistic motion for much longer and their velocities remain more correlated over time (Fig. S4B). The diffusive motion of osteoblasts within the differentiated, more densely packed (S2B-D), collagen-rich bone center is consistent with the idea that these cells experience a different stiffness from those at the softer end of the tissue. Such MSD patterns suggest that material properties such as stiffness influences features of cell motion, and in particular, restricts a cell’s ability to move persistently. Inhomogenous tissue level forces similar to ones observed here are known to regulate proliferation rates and division orientation in other systems (*19, 20*). We found reduced proliferation within the bone center which harbors the most collagen and cells are more densely packed, consistent with inhibited proliferation in densely packed environments (Fig. S2E; S3B, C) (*21*). We also found that labeled osteoblasts divided along the axis of growth in both live imaging and fixed skull caps (Fig. 1J-L; Supplemental Movie 3). Not only were the medial-lateral relationships between daughter cells as well as neighbors maintained for the duration of the movie (Fig. 1M, N; Supplemental Movie 4) but daughter cell displacement was greatest toward the front (Fig. 1O, P). Together these data support a role for stiffness and collagen gradients in regulating cell dynamics during bone morphogenesis where the bone front is increasingly compliant when compared to the bone center.

### Inhomogenous structure is built by progressive osteoblast differentiation ahead of the bone

While tracking individual nuclei we noticed that cells initially at the osteogenic front, which separates differentiated from undifferentiated cells, were no longer at the leading edge by the end of our live imaging. To confirm this finding we quantified the relative displacement of the front and compared this displacement to that of individual cells originally residing at the front. We found displacement of the osteogenic front to be consistently greater than that of tracked nuclei (Fig. 2A-C). While we found the expansion rate of the osteogenic front ((15 ± 6) μm) to be comparable to *in vivo* measurements between E13.5 and E14.5 (Fig. S1D) ((25 ± 4) μm), tracked nuclei moved more slowly compared to the front interface (Fig. 2C). These data suggest that newly differentiated osteoblasts are added to the osteogenic front as the motion of tracked cells cannot explain the extension of the front.

**Figure 2.**
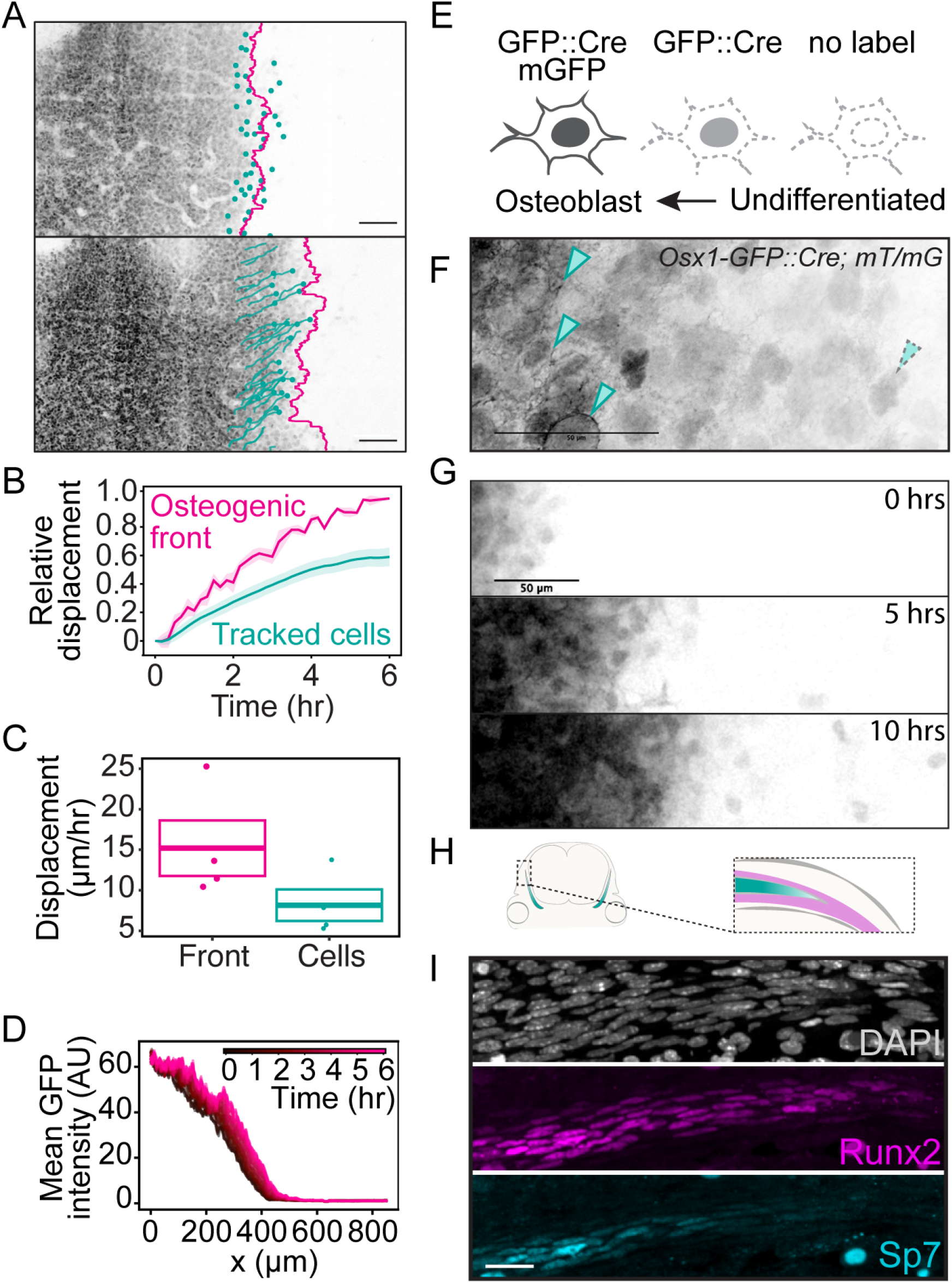
*Ex vivo* live imaging of bone expansion reveals progressive differentiation of osteoblasts. **(A)** Max projection of *Osx1-GFP::Cre* labelled frontal bone at 0 h and 6 h together with the osteogenic front (magenta) and individually tracked cells (cyan). Scale bar = 100 mm. **(B)** The relative displacement for the osteogenic front (magenta) and tracked cells (cyan) over the course of 6 h, defined as the displacement normalised by the total displacement of the osteogenic front at 6 h. Shaded areas show SEMs (N=4). **(C)** Rate of expansion at the osteogenic front compared to that of tracked cells. **(D)** Average GFP intensity profiles along the medial-lateral axis for a labelled frontal bone in an *ex vivo* imaging experiment starting at E13.75. Each graph arises from one time frame, with the color indicated in the legend. **(E)** GFP labelling scheme in *Osx1-GFP::Cre; R26RmT/mG* explants. **(F)** Representative fixed tissue image of *Osx1-GFP::Cre; R26RmT/mG*. **(G)** Max projections showing GFP localization in E13.75 live movies of *Osx1-GFP::Cre; R26RmT/mG* explants at 5 h time intervals. Scale bars = 50 mm. **(H)** Schematic of a coronal section with the bone labelled in cyan. The inset shows the imaged areas of I with the approximate domains of Sp7+ osteoblasts (cyan) and Runx2+ precursor cells (magenta). **(I)** DAPI, Runx2 and SP7 immunoreactivity at osteogenic front in E14.5 coronal sections.

Additional osteoblasts at the osteogenic front could arise from the differentiation of mesenchyme that resides medial to the developing bone. If so, we would expect to find greater GFP intensity in osteoblasts in the bone center compared to those at the front as GFP signal increases with osteoblast maturation in *Osx1-GFP::Cre* reporter mice. To test this, we quantified GFP intensity in osteoblasts across the mediolateral axis of the bone and indeed found nuclei decreased in signal intensity toward the front (Fig. 2D). Further, this intensity profile was shifted over time indicating an increase in GFP expression that could arise from newly differentiated cells. To confirm that cells newly expressing *GFP* have only recently activated the *Osx1* promotor and have not lost and reactivated *Osx* driven expression, we crossed the *Osx1-GFP::Cre* line to *R262RmT/mG* (*22*) reporter mice. Here, membrane GFP would be driven after accumulation of Cre recombinase in our *Osx1* reporter and thus newly differentiated osteoblasts would harbor only nuclear GFP whereas membrane GFP would be expressed in addition to nuclear GFP in maturing osteoblasts (Fig. 2E-F). Indeed, we found nuclear-only signal at the osteogenic front where osteoblasts in the bulk of the bone were labeled with both membrane and nuclear GFP (Fig. 2G; Supplemental Video 5). Likewise, we found expression of an early skeletal marker, *Runx2* to be expressed ahead of the osteogenic front in *Osx1* negative mesenchyme (Fig. 2H-I). These data demonstrate that differentiation occurs at the osteogenic front and that cell motion is insufficient to explain the dynamics of bone growth.

### Feedback between tissue material properties and cell fate is sufficient to drive noncanonical antidurotaxis

We next asked whether coupling between tissue material properties and cell fate is sufficient to drive key features of calvarial morphogenesis as observed. To this end, we constructed a minimal theoretical model describing the tissue as a viscous fluid with two types of cells (osteoblasts and undifferentiated mesenchyme) which undergo effective proliferation and differentiation and are subject to cellular flows arising from force imbalances (Fig. 3A; Supplementary Text). We modeled differentiated osteoblasts to generate a stiffer environment than undifferentiated mesenchyme to reflect differential rates of collagen production. Conversely, we modeled a differentiation rate that increases with higher stiffness, in accordance with *in vitro* experiments (*23, 24, 24–30*), thus generating a mechanical feedback loop between cell fate and stiffness.

**Figure 3.**
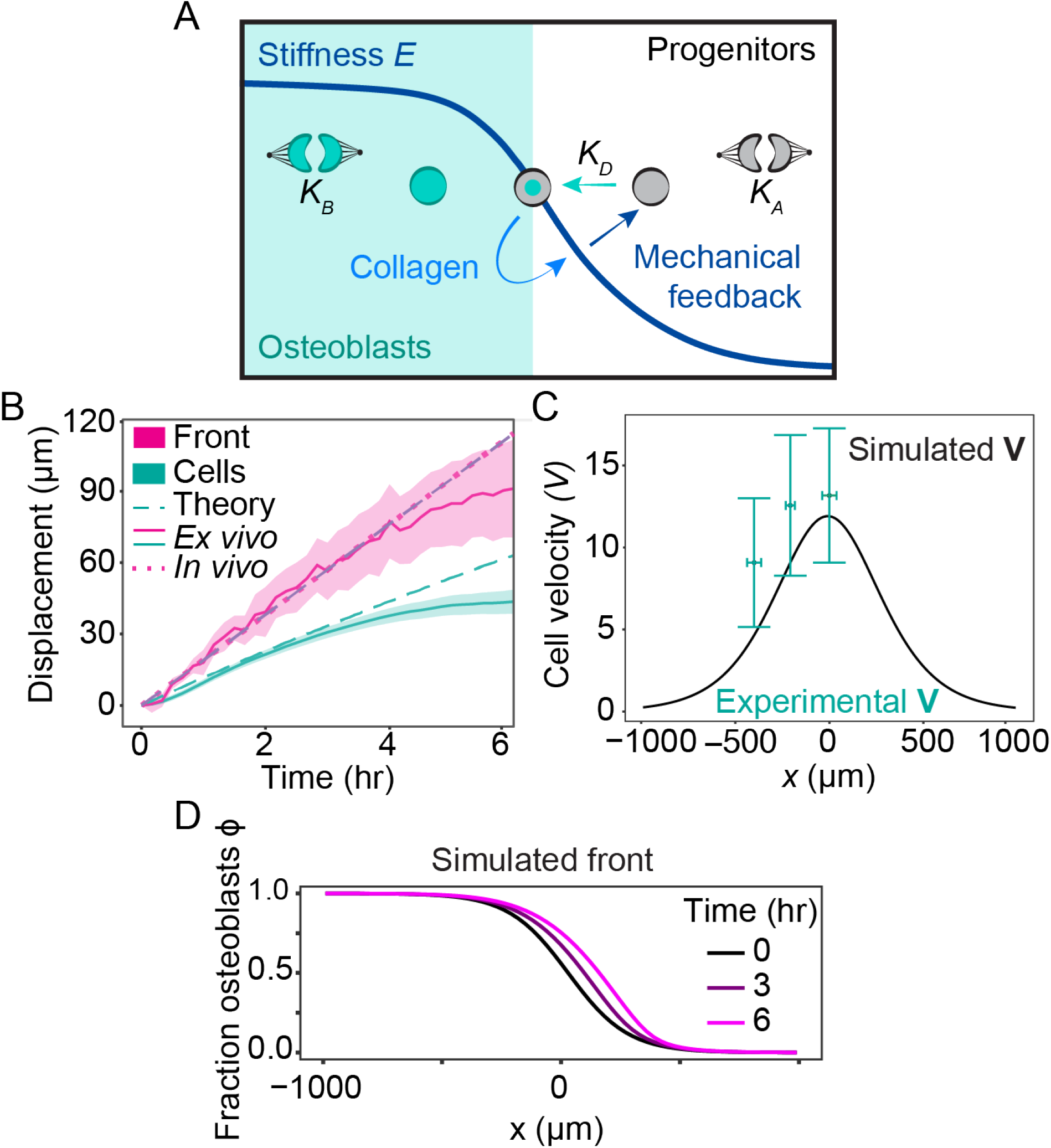
Biophysical model with mechanical feedback between stiffness and cell fate recapitulates imaging results. **(A)** Schematic of the model. **(B)** Dynamics of the osteogenic front and the tracked cells for the theoretical model, the *ex vivo* live imaging experiments (N=4) and *in vivo* fixed tissue images obtained by comparing bone sizes between E13.5 (n=7) and E14.0 (n=7). Shaded areas indicate SEM for the *ex vivo* data. **(C)** Cell velocity for the theoretical model compared to the velocities obtained from the *ex vivo* tracked cells over 6 hours. Error bars show standard deviations. **(D)** The simulated fraction of osteoblasts shows a sigmoidal profile across space that travels as a wave. The horizontal axis shows the position along the medial-lateral axis, where the origin represents the initial position of the osteogenic front, here defined to be the location where *ϕ* = 1*/*2.

We simulated the model with realistic biological parameters, including those quantified in this study. Our model generates an expansion of the osteoblast domain with differential velocities for the osteogenic front and for individually tracked cells, which can be simultaneously fit to the experimentally measured values (Fig. 3B; Fig. S5). Moreover, upon fitting only these expansion velocities, the model correctly predicted the spatial profiles of the relative osteoblast concentration (corresponding to GFP intensity) and cell velocity. Specifically, the cell velocity profile shows a peak at the osteogenic front with cells near the front moving faster than cells away from the front (Fig. 3C).

Our model also predicted the observation that the spatial profile of osteoblast density is characterized by a stable, expanding wavefront of differentiated osteoblasts (Fig. 3D), as found by quantifying the measured GFP intensity profiles (Fig. 2D). Although such a wavefront may arise in tissues with spatially inhomogeneous cell division rates (*31*), our PH3 immunostaining revealed no significant differences in proliferation rates between the bone front and undifferentiated mesenchyme (Fig. S2E). Therefore, we proposed that the osteogenic wave is instead driven by the aforementioned mechanical feedback (Supplementary Text). Further quantifications of fluctuations of the osteogenic front confirm that the data are consistent with a biophysical wave with mathematical properties described by our model (Fig. S4C-F; Supplementary Text). Altogether, these data support the idea that a biophysical wave generated by mechanical feedback is sufficient to recapitulate complex tissue dynamics during skull morphogenesis.

### Perturbing the stiffness gradient changes bone size

As our model predicts that a stiffness gradient is sufficient to drive both cell motion and differentiation toward the midline, we hypothesized that altering the stiffness gradient should perturb bone expansion. To test this we perturbed the structure of the emergent collagen network by inhibiting Lysl Oxidase (LOX), a key collagen crosslinker (Fig. 4A-C). We found that Beta-Aminoproprionitrile (BAPN), an irreversible competitive inhibitor of LOX (*32–36*) reduced tissue stiffness at the osteogenic front to generate a greater stiffness gradient when compared to control samples (Fig. 4D). Our model predicts that a greater difference in tissue stiffness between the bone center and front would promote the expansion of frontal bones. Indeed, we found significantly larger frontal bones toward the end of skull expansion (Fig. 4E-G). These data are consistent with the predictions of our model and suggest that a gradient in tissue stiffness or pressure generated by differentiation-dependent collagen production is sufficient to orchestrate frontal bone morphogenesis (Fig. 4H).

**Figure 4.**
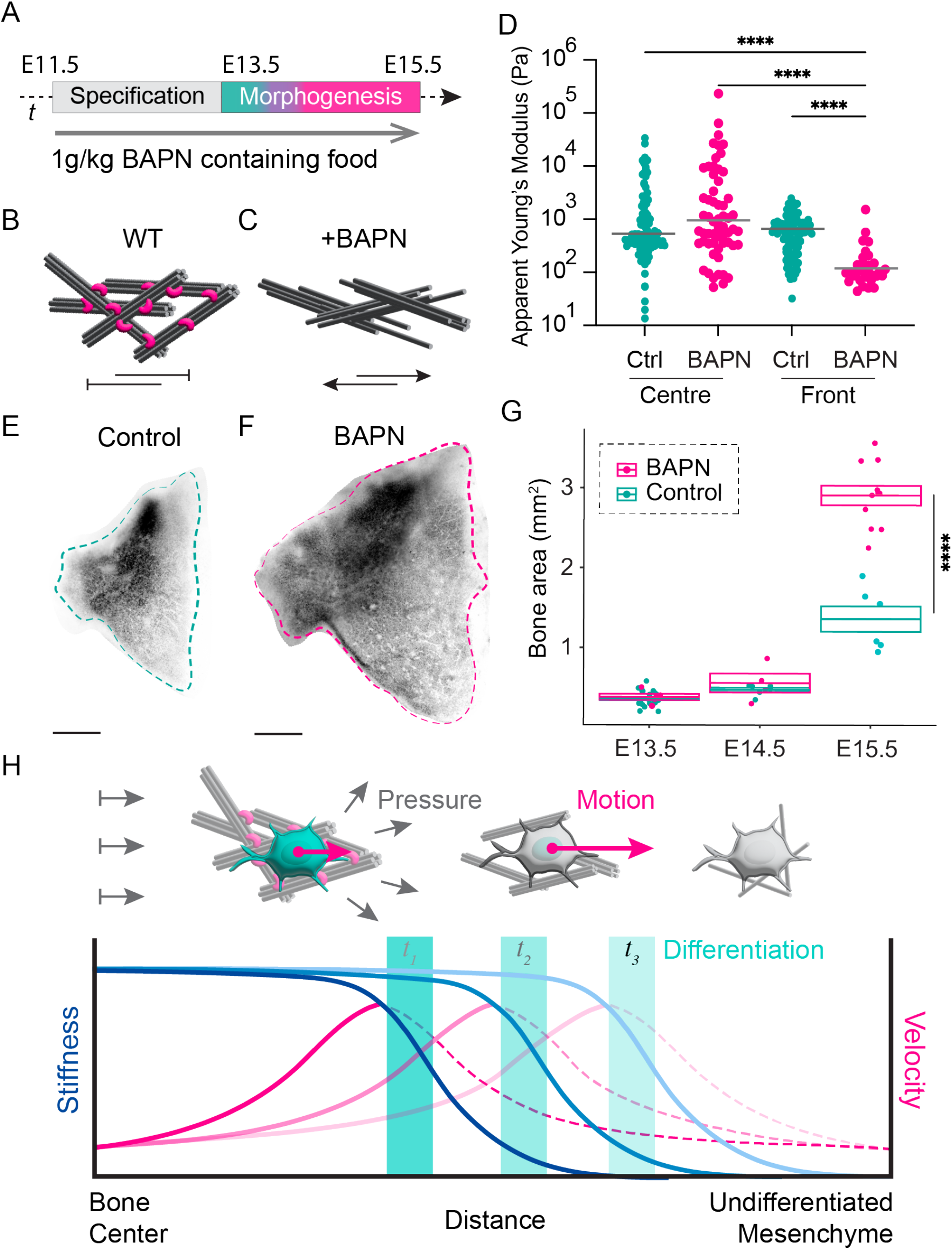
Perturbing collagen crosslinking leads to a larger difference in stiffness across the tissue resulting in larger bones. **(A)** Schematic of the BAPN feeding protocol. **(B)** Schematic showing collagen (black) fibrils crosslinked by Lysyl Oxidase (magenta). **(C)** Schematic showing collagen without crosslinkers in BAPN treated embryos. **(D)** AFM measurements of the stiffness (apparent Young’s Modulus) at E15.5 at two different locations of the tissue, for control samples (cyan) and BAPN-treated samples (magenta), with horizontal lines indicating the median (Kruskal–Wallis test, *p <* 0.0001). **(E)** Representative image of a control *Osx1-GFP::Cre* labelled frontal bone at E15.5. **(F)** Representative image of a BAPN-treated *Osx1-GFP::Cre* labelled frontal bone at E15.5. **(G)** Box plot together with individual data points showing measured frontal bone areas for the two conditions at E13.5, E14.5, and E15.5 (Mann-Whitney test, *p <* 0.0001). **(H)** Schematic showing a proposed model of anisotropic frontal bone expansion.

## Discussion

We discover an alternate model of motility that does not depend upon the intrinsic dynamics of individual cells. Although stiffness gradients are known to drive durotaxis *in vivo* (*37*) as well as anti-durotaxis *in vitro* (*38*), it has not been shown that such gradients can generate motion without requiring active migration. We show how self-organized mechanochemical coupling between stiffness and cell fate drives cell motility, oriented cell division, and cell differentiation to ultimately control tissue size. Our work demonstrates a general mechanism for generating anisotropic tissue growth during skull morphogenesis and in mesenchymal tissues more broadly. Understanding mesenchymal collective cell motility could improve our understanding of other processes involving mesenchymal cells such as tissue homeostasis, stem cell maintenance, and blood vessel structure, in both healthy and cancerous states (*39*).

Although we are limited to analyzing nuclear dynamics and cannot infer the dynamics of cell borders, tracking these nuclei revealed that motion is not equal across the bone primordia. While both cell flows and migration could give rise to similar spatial dynamics, we find little evidence of active cell migration. First, it is important to note that osteoblasts are not moving on a substrate. Rather, they are embedded in the tissue and do not directly interact with the epithelial basement membrane or fibrillar matrix of the meninges. We also find cellular motion to be down the stiffness and collagen gradient, whereas durotaxis most commonly drives migration up these gradients of mesenchymal cells and the tissue composition is not consistent with fibroblastic anti-durotaxis (*3, 40*). Further, there are no true leader cells and cell rearrangements are very limited, contrary to what is typical during collective cell migration (*1*). Our data can be explained without the need for collective cell migration which is contrary to previous assumptions (*15, 16*). Instead, our data indicate that differentiation plays a predominant role in establishing the inhomogenous physical environment that also generates motility.

While previous *in vitro* studies have found that osteoblast differentiation increases with substrate stiffness (*41,42*), the stiffness values measured in our study are orders of magnitudes lower than is typically required for osteoblast differentiation *in vitro*. This highlights the need for *in vivo* studies to confirm mechanisms found *in vitro* and to establish physiological parameters in developing tissues, which cannot always be directly extrapolated from these *in vitro* studies.

Our perturbation experiments also demonstrated that the slope of the stiffness gradient determines bone size, in accordance with our biophysical theory. The stiffness gradient slope could therefore function as a tunable parameter controlling overall skull size and enable coexistence of a variety of skull sizes in different organisms. Fundamentally, our work demonstrates that conceptual mechanisms driving molecular motion in non-living physical systems also extend to cell motility at the scale of tissues.

Together, this work extends physical principles which regulate dynamics at the molecular and subcellular scales to that of tissues. We provide a new conceptual framework to understand morphogenesis and motility in biology that emerges within collectives. In asking whether cellular mechanisms such as durotaxis found *in vitro*, hold true in tissues, it is essential that we consider noncanonical forms of motility. Understanding alternative modes of motility has wider implications for engineering tissues *in vitro* and interrogating mechanisms of morphogenesis *in vivo*.

## Acknowledgements

We thank the Biomedical Services Facility, the Light Microscopy Facility and the Electron Microscopy Facility at the MPI-CBG, for assistance with data collection and optimization of imaging protocols. We thank members of the Tabler Lab, Stephan Grill, Michele Marasss, Anais Bailles, Ricard Alert, Marko Brankatschk, and Jan Brugues for helpful discussions and critical feedback. This work was funded in part by the DFG (TA1515/1-1). YD was funded in part by an Elbe Fellowship (MPI-CBG and CSBD).

## Author Contributions

Conceptualization: J.M.T. Methodology: J.M.T, Y.D. A.T, and E.U. Investigation: J.M.T, Y.D., A.L.C, D.A.A, J.L, A.T, and E.U. Visualization: J.M.T and Y.D. Supervision: J.M.T and S.R. Writing: J.M.T and Y.D.

## Data and materials availability

The authors confirm that the data supporting the findings of this study are available within the article and its supplementary materials.

## Supplementary materials

Materials and Methods

Supplementary Text

Supplementary Figures S1 to S5

Supplementary Movies S1 to S5

## Materials and Methods

### Mouse lines

The following mouse lines were used: *Sp7-mCherry* (Tg(Sp7/mCherry)2Pmay/J, available via Jax) (*43*),

*Osx1-GFP::Cre* (B6.Cg-Tg(Sp7-tTA,tetO-EGFP/cre)1AMC/J, available via JAX) (*18*), *R26R-mT/mG* (GT(Rosa)26Sortm4(ACTB-tdTomato-EGFP)Luo, available via JAX) (*22*) and C57Bl/6JOlaHsd or C57Bl/6NTac. Genotyping was performed as described in original publications. Experiments conducted at the University of Texas, Austin were performed in accordance with approved IACUC protocols. Experiments conducted at the MPI-CBG adhered to the German Welfare Act and were overseen by the Institutional Animal Welfare Officer.

### Lysyl-Oxidase Inhibition

For collagen crosslinking inhibition studies, pregnant *Osx1-GFP::Cre* females were fed 0.25% (1 g*/*kg) BAPN containing food from E11.5, the onset of skull morphogenesis. For embryo collection, pregnant females were euthanized by cervical dislocation and embryos were collected for downstream analysis. Protocol was approved by the Institutional Animal Welfare Officer. BAPN containing food made by Safe Complete Care Competence.

### Culture Media

High Glucose DMEM (Sigma D6469) is supplemented with 10% Fetal Calf Serum (Gibco A3160502), 100 mg*/*mL ascorbic acid (Sigma, PHR1008), 10 μM beta-glycerolphosphate (Sigma 50020), 0.5 mL in 50 mL 100 X Antibiotic Antimycotic (Sigma 15240062).

### Imaging chamber preparation

Sarstedt Lumox dish 35 chambers were coated in Fibronectin from bovine plasma (Sigma F1141-1MG) diluted 1:1 with DMEM. (Sigma D6469). Dishes were left to dry (∼ 15 minutes) while embryos, and media were prepared. A cyclopore polycarbonate membrane filter (GE Healthcare Whatman, 7060-2516) and silicone well (Flexiperm ConA, Sarstedt, 96077434) for holding the sample in place were placed in DMEM after being washed in 70% Ethanol in advance of sample preparation.

### Immunohistochemistry

All immunohistochemistry stainings were performed according to standard protocols. All embryos were collected in cold PBS and fixed in 4% PFA. After fixation embryos were embedded in 15% sucrose/7.5% gelatin and frozen in dry ice. 35 μm coronal sections were collected for nuclear shape and nuclear envelope analysis. Cryosections were stained with Lamin A/C, counterstained with DAPI (1:500, Sigma, 32670) and then coverslipped with Vectashield. For immunohistochemistry antibody staining 20 μm coronal sections were collected. Antigen retrieval was performed in 10 nm sodium citrate (pH 6.0). Sections were blocked with 10% goat serum in PBS for 1 hour at room temperature and incubated with primary antibodies overnight at room temperature. A list of the used primary antibodies can be found on Table 1. Coronal sections were then incubated with secondary antibodies for 2 hours at room temperature, counterstained with DAPI and then coverslipped with Vectashield (Vector Labs, H1000).

**Table 1:** Antibodies and dilutions.

| Antibody | Manufacturer | Catalog Number | Dilution |
| --- | --- | --- | --- |
| Mouse anti-Lamin A/C | Cell Signalling | 8617S | 1:50 |
| Mouse anti-Twist2 | Abcam | ab50887 | 1:50 |
| Mouse anti-Runx2 | Santa Cruz Biotechnology | sc-390715 | 1:50 |
| Alexa Fluor 488 Goat anti-Mouse IgG | Invitrogen | A-11001 | 1:500 |
| Alexa Fluor 568 Goat anti-Rabbit IgG | Invitrogen | A-11036 | 1:500 |

**Table 2:** Estimation of model parameters from experiments, literature and parameter fitting.

| Symbol | Interpretation | Estimate | Source |
| --- | --- | --- | --- |
| $\rho_A^c$ | Carrying capacity A cells | $5.8 \cdot 10^{-3} \mu m^{-2}$ | Fixed tissues imaging (Fig. S2D) |
| $\rho_B^c$ | Carrying capacity B cells | $7.6 \cdot 10^{-3} \mu m^{-2}$ | Fixed tissues imaging (Fig. S2D) |
| $D$ | Diffusion constant | $15 \mu m^2/hr$ | Live imaging cell tracks (Fig. S2B) |
| $E_M$ | Stiffness cell type A | 30 Pa | Nanoindenter measurements (Fig. S2A) |
| $E_O$ | Stiffness cell type B | 1000 Pa | Nanoindenter measurements (Fig. S2A) |
| $\tau$ | Timescale in Eq. 8 | 10 hr | Estimate from (55) |
| $\eta$ | Viscosity | $10^4 Pa \cdot s$ | Typically $10^3 - 10^5 Pa \cdot s$ (52, 60, 61) |
| $\gamma$ | Friction coefficient | $0.8 \times 10^3$ | Fit cell velocity; typically $10^3 - 10^5 \frac{Pa \cdot s}{\mu m^2}$ ((61, 62)) |
| $\alpha$ | Coefficient in Eq. 18 | $10^{-6} Pa^{-1} s^{-1}$ | Fit front velocity $v_F$ using Eq. 30 |

### Embryo manipulation for osteoblast live imaging

E13.75 embryos are extracted from yolk sacs and scored for reporter gene expression in DMEM (Sigma D6469). Skull caps comprising epidermis, the paired frontal and parietal bones, and meninges were excised with Vannas spring scissors 2.5 mm (F.S.T,15001-08), as indicated in Fig. 1[B], in fresh DMEM. The skull explant was transferred to the center of the pre-prepared gas permeable dish, meningeal (basal) side-down with the convex side of a 130 mm double spatula. DMEM washed membrane, is dried slightly by dabbing it once on tissue paper. The membrane was then gently laid down over the sample, being careful to avoid bubbles. Next, the washed silicone well was dabbed dry with tissue paper and quickly placed on top of the permeable membrane such that the sample sits underneath the center of the well. The dish was then filled with 2.5 mL culture media, followed by a thin layer of mineral oil (Sigma M-8410). The sample was then transferred to the environmental chamber of the microscope.

### Flat-mount skull cap imaging of fixed samples

After excising and fixing, skull caps were mounted flat in vectashield and imaged using the Zeiss AxioZoom ApoTome system.

### Second Harmonic Generation

*Osx1::GFP-Cre* skull caps were excised and mounted as before and imaged using 2-photon excitation system of the Leica DMI 4000 and Olympus UplanSApo 40x/0.90 Dry objective. 488 laser lines were used to excite the GFP::Cre fusion protein and 950 nm for second harmonic generation. We also collected simultaneous images of Nucelar auto fluorescence at 490 nm using 950 nm excitation.

### Life-Act viral reporter infection

Skull caps were excised as before and placed in a 2 mL microcentrifuge tube for 1 hour in 50 μL of 106 IU adenoviral IrAVCMV-LifeAct-TagRFP construct (1×1010 IU/ml, Ibidi, 60122) diluted in culture media. Adenoviral solution was then removed and skull caps were washed twice in culture media. Adenoviral contaminated media and tips are incubated in 10% bleach for 20mins according to BSl2 (USA) and S2 (EU) regulations. Explants were then plated as above and incubated at 37 μm in a humidified incubator for 18 hours and then transferred to the microscope and imaged.

### Live imaging

Our explant system is adaptable to almost any inverted confocal microscope with an air objective and resonant scanner. Three different confocal microscopes were used in this study. Movies were captured on the single photon Nikon TiE system with CFI60 Plan Apochromat Lambda 40x Objective Lens. Next, we used a 2-photon excitation system of the Leica DMI 4000 and Olympus UplanSApo 40x/0.90 Dry objective. 488 laser lines were used to excite the GFP::Cre fusion protein. Later, explants were imaged Andor Revolution WD Borealis Mosaic (Andor) Spinning Disk confocal using the Olympus UplanSApo 40x/0.90 Dry objective.

### Live imaging stitching

To facilitate downstream analysis of multiple position acquisition time lapse imaging data, we used an ImageJ/Fiji (*44, 45*) script developed by the Max Planck Institute for Cell Biology and Genetics Scientific Computing Facility (https://github.com/PreibischLab/BigStitcher/). The script converts each tile of the raw multiphoton imaging data set into the tiff image format then uses the BigStitcher plugin for ImageJ/Fiji to stitch the tiles (*46*).

### Manual Live Imaging Analysis

To facilitate image analysis using tools available in the base ImageJ/Fiji package, a maximum intensity projection of live imaging was generated using the Z-projection plugin. To obtain *Osx1-GFP::Cre* intensity measurements along the medial lateral axis, we projected and averaged the intensity of these maximum intensity projected images over the vertical axis.

### Atomic Force Microscopy

For measurements of tissue bulk stiffness embryos were collected in cold 1x PBS, heads were dissected and embedded in 4% low gelling agarose (Sigma, A4018). 2 mm sections were obtained using a vibratome (Leica, VT1200S) and immobilized using tissue seal (histoacryl blue) in a polystyrene-bottom dish (TPP, 93060). Measurements were performed using a Cellhesion 200 (JPK Instruments/Bruker) mounted on top of a Zeiss Axio Zoom (Zeiss, .V16). The cantilevers (arrow T1, NanoWorld), modified with 20 μm diameter polystyrene beads (microparticles GmbH), were calibrated by the thermal noise method using built-in procedures of the SPM software (JPK Instruments). Measurements were performed at room temperature (18 °C-20 °C). Individual force-distance curves were acquired with defined approach and retract velocity (7.5 μm*/*sec) and with a contact forces ranging from 2.5 nN-10 nN in order to reach approximately constant indentation depths of 2 μm. At least five specimens were probed for each tissue in a 25 μm grid at 5 μm intervals. The apparent Young’s modulus *E* was extracted from approach force–distance curves by fitting to the Hertz/Sneddon (*47, 48*) model for a spherical indenter using JPK data processing software.

### NanoIndentation

Nanoindentation was performed using the Chiaro Nanoindentor system (S-Chairo-ST, third generation). Samples were plated on a gas permeable membrane and for measurements, media was removed. Samples were viewed using Zeiss Inverted CCD and 5x/0.15 Plan-Neofluar, Air, Ph1, Zeiss objective.

### Cell tracing and quantification of cell tracks

The Manual Tracking plugin in ImageJ/Fiji was used to record the motion of osteoblasts at different positions in the bone. Tracks were recorded for osteoblasts at three different positions across the bone: 1) osteoblasts at the osteogenic front 2) osteoblasts approximately 200 μm lateral to the front, and 3) osteoblasts approximately 400 μm lateral to the front.

From the cell tracing we obtained individual cell tracks consisting of sets of two-dimensional positions 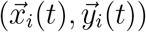 for 1 ≤ *i* ≤ *N* at different times 0 ≤ *t* ≤ *T*. From this we directly computed the average cell velocities 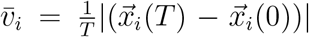 and mean squared displacements 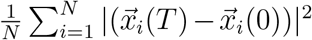. To compare directions of motion, we computed a spatial correlation function between normalized velocities of cells, defined as

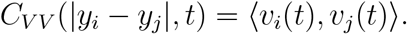

Here *y*_*i*_, *y*_*j*_ indicate the y-positions of cells *i* and *j, v*_*i*_(*t*), *v*_*j*_(*t*) their normalized velocities at time t, and the brackets indicate averages over all cells at a distance |*y*_*i*_ − *y*_*j*_| from each other (in practice obtained from binning cells with comparable distances to each other).

### Division orientation

Divisions were identified by observing metaphase plate formation and anaphase on subsequent time frames. Using the centroids of each daughter cell the angle of divisions was determined using the ImageJ/Fiji ImageJ/Fiji angle tool relative to the angle of the osteogenic front. In this measurement scheme 90° represents a cell dividing parallel to the medio-lateral axis of expansion.

### Daughter displacement

The distance that each daughter cell displaced along the axis of expansion was measured using the ImageJ/Fiji line tool. The centroid of each daughter cell was measured relative to the position of the metaphase plate centroid along the axis of expansion.

### BAPN protocol

1 g*/*kg (Sigma Aldrich, A3134) containing food was fed to pregnant dams from E11.5 until collection at E15.5.

### Quantifying bone sizes (fixed tissue images)

We obtained fixed tissue images of the *Osx1::GFP-Cre* and *Sp7-mCherry* lines for WT and BAPN at equal magnifications for E13.5, E14.5 and E15.5. We first segmented the images to extract the region corresponding to the frontal bone in each image. We did this by thresholding the images using a common intensity threshold, and then selected the largest connected region corresponding to the frontal bone for each image.

## Supplementary Text

### Theoretical model

We constructed a continuum model for a tissue with two cell types, described in terms of spatially and temporally varying fields of cell density, cell flows and mechanical stresses. Let *ρ*_*i*_(*x, t*) (*i* = *A, B*) be the concentrations of cells of type A and B. Although the model formulation is generic, in the context of our system we take *A* to be the undifferentiated mesenchymal progenitors and *B* the differentiated osteoblast population. Consider two different processes that can change these concentrations:

1. Proliferation and cell death give rise to effective reproduction rates *k*_*i*_(*ρ*_*i*_) (*i* = *A, B*), which depend on local cell densities as described below.
2. *A* is converted to *B* irreversibly at a rate *k*_*D*_, which can depend on other (dynamic) variables of the system, as will be discussed in the next section.

Number balance imposes the following dynamic equations on the cell densities *ρ*_*A*_, *ρ*_*B*_:

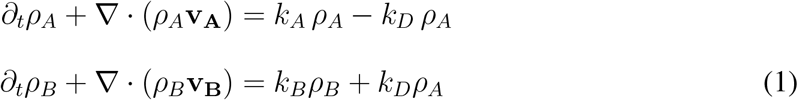

These equations describe how the local density of *A* and *B* cells can change through advection (with velocities *v*_*A*_, *v*_*B*_), cell division or loss, as well as conversion of *A* cells into *B* cells. As we are interested in the differentiation dynamics of osteoblasts, we make the following transformation (*31*):

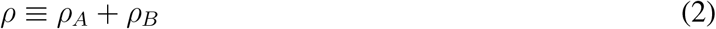

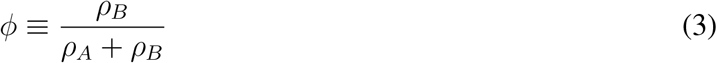

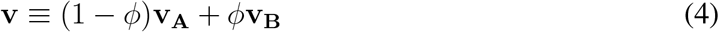

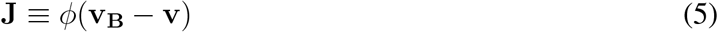

Hence, rather than considering individual densities *ρ*_*A*_, *ρ*_*B*_, we describe the system in terms of the total concentration of cells *ρ* and the local fraction of *B* cells *ϕ*. Similarly, rather than considering individual cell velocities **v**_**A**_, **v**_**B**_, we consider the **v, J**. If we assume that the mass per cell of both cell types is roughly equal, then **v** represents the center of mass velocity and **J** represents the relative flux of *B* cells with respect to the center of mass.

From Eq. 1, we then derive the following equations for *ρ*(*x, t*) and *ϕ*(*x, t*):

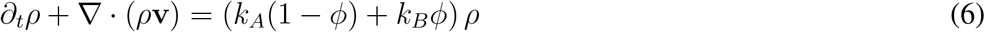

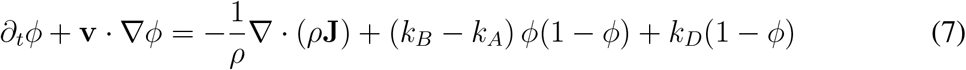

We assume **J** = −*D*∇*ϕ*, in analogy to Fick’s law for diffusion, but with respect to the relative fraction of *B* cells *ϕ* rather than total concentration. Hence we assume that in the absence of other processes, local differences in cell type composition will eventually be relaxed to a homogeneous state where the local composition is uniform across space.

We impose constraints on tissue growth by assuming that the cell density essentially follows a logistic growth model, such that the net growth rate decreases when the density becomes too high (*49, 50*). Therefore, we write the net division rate as

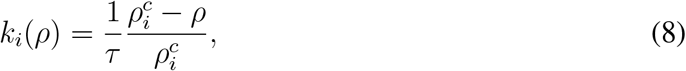

where 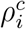 with *i* = *A, B* are the carrying capacities for the two cell types and *τ* sets the time scale of adapting to homeostasis.

To solve the above system of PDEs, we need an additional equation for the advection velocity **v**, which we derive from force balance. Various studies have argued that the material properties of tissues largely follow that of a viscoelastic medium, with the property that at sufficiently long timescales we can ignore elastic terms and the tissue effectively behaves as a viscous fluid (*11, 50–52*). If we assume the medium is isotropic, the stress tensor in the tissue can be decomposed as (*53*)

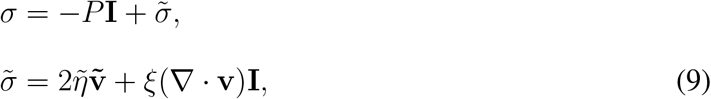

where *P* is the pressure, **I** is the identity matrix, 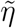 is the dynamic viscosity, *ξ* is the bulk viscosity, 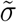 is the deviatoric stress tensor and its traceless part 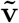 is defined as

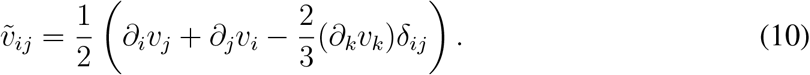

Force balance in the tissue then takes the general form

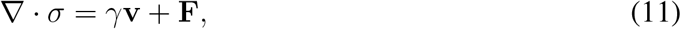

where *γ* is the friction coefficient and **F** represents external forces acting on the tissue.

Since the tissue we study consists of a thin sheet of cells, we will describe it as an effective 2D system and neglect the *z* dimension by averaging all quantities in this direction. Furthermore, as our data shows that cells persistently move unidirectionally and that this directionality is correlated over large distances in the direction perpendicular to that of cell motion (Fig. S4A), so to first approximation we can set *v*_*y*_ = 0 and *∂*_*y*_*v*_*x*_ = 0, such that we obtain an effective one-dimensional description of our system. In particular, using Eqs. 9 and 11 and writing 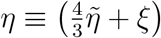 and in the absence of external forces (**F** = 0) we now obtain

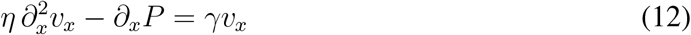

To close the system and solve for *v*, we require an equation of state of the form *P* (*ρ*). For tissues, the following form derived from thermodynamics has been proposed (*54, 55*):

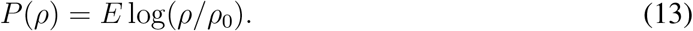

Here *E* is the Young’s modulus, which in general can be spatially dependent (as will be discussed below), and *ρ*_0_ sets the density at which the pressure is defined to be zero.

### Mechanical feedback

To capture the interaction between material properties and cellular processes in the above system, we introduced a feedback loop in the model that couples these two features. First, we modelled the stiffness to depend locally on the fraction of differentiated osteoblasts through

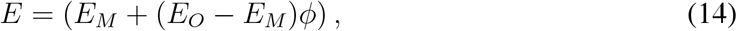

where *E*_*M*_, *E*_*O*_ represent the stiffnesses generated by populations consisting of only mesenchymal and osteoblast cells respectively. This reflects the fact that osteoblasts mineralise the ECM in their immediate vicinity and generate a stiffer environment than mesenchyme progenitors, as shown in our stiffness measurements.

Second, we let the differentiation rate *k*_*D*_ = *k*_*D*_(*E*) depend on the local stiffness *E*, in accordance with *in vitro* experiments establishing a clear role for substrate stiffness in regulating osteoblast differentiation (*23,24,24–30*). In particular, stretch and compression of tissues and/or substrate have been shown to increase osteoblast differentiation and it has been suggested that this is mediated through activation of BMP signaling, ERK signaling and p38 MAPK phosphorylation (*56*). Although the exact quantitative relation between stiffness and differentiation is not known *in vivo*, we next derive a few constraints on this relation based on the property that the osteoblast expansion dynamics follows that of a linear unstable wave.

### Wave solutions

As the osteoblast expansion is consistent with that of an unstable linear wave, also known as Fisher-Kolmogorov-Petrovsky–Piskunov (FKPP) wave (*57, 58*) and the mathematical conditions under which such waves can arise are well-known (*59*), we can now analyse conditions under which such a wave is generated in our model. We identify two possible mechanisms capable of generating a Fisher-KPP wave in our system: (1) a difference in proliferation rates *k*_*B*_ −*k*_*A*_ *>* 0, or (2) a non-linearity in the term describing differentiation from *A* to *B, k*_*D*_(1−*ϕ*). To see this, note that Eq. 27 in the absence of density gradients (*∂*_*x*_*ρ* = 0) and cell movement (*v* = 0) reduces to a generalized FKPP equation of the form

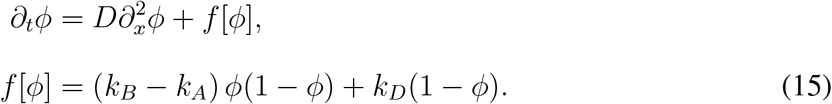

In our system, the first scenario of a proliferation gradient would imply that the osteoblasts would divide more than the mesenchymal progenitors and generate a pressure gradient that pushes progenitor cells towards the lateral side of the tissue (*31*). However, our PH3 immunostaining revealed no significant difference between the bone front and undifferentiated mesenchyme in terms of proliferation rates (*k*_*A*_ ≈ *k*_*B*_). Consequently, we consider the second scenario and set *k*_*B*_ − *k*_*A*_ = 0, so we have *f* [*ϕ*] = *k*_*D*_(1 − *ϕ*), where *k*_*D*_ is an yet unspecified term depending on the stiffness *E* and subsequently through Eq. 14 on *ϕ*. The general condition for an FKPP wave (*59*) then implies that *f* [*ϕ* = 0] = *f* [*ϕ* = 1] = 0, while *f*^*′*^[*ϕ* = 0] *>* 0 and *f*^*′*^[*ϕ* = 1] *<* 0. Translated into conditions for *k*_*D*_, we first get the condition

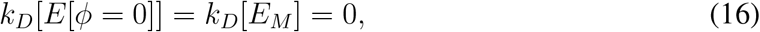

implying that the differentiation rate should be zero when the stiffness of the system is that of mesenchymal cells, i.e. in the absence of osteoblasts. Secondly, the conditions on the derivatives of *f* [*ϕ*] imply

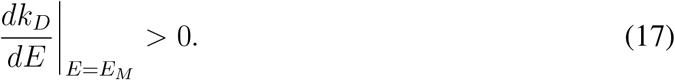

This condition can be interpreted as saying that any increase in stiffness from the mesenchymal value *E*_*M*_ should lead to an increase in *k*_*D*_, and therefore a positive differentiation rate. In other words, while strictly at the mesenchymal stiffness the differentiation rate should be zero, any slight increase would tilt the differentiation rate constant toward positive values.

In the absence of detailed knowledge about the functional form of *k*_*D*_, in our simulations we assume a simple linear relation:

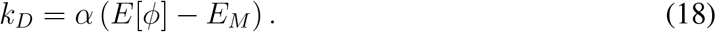

Together with Eq. 14, this implies that the total differentiation rate takes the form corresponding exactly to the classical FKPP equation

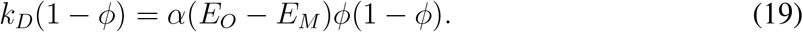

In this picture, the differentiation rate constant *k*_*D*_ increases monotonically with stiffness, but the differentiation rate *k*_*D*_(1 − *ϕ*) shows a single maximum at *ϕ* = 1*/*2. This is due to the depletion of progenitor cells as *ϕ* and thereby the stiffness increases, which therefore reduce differentiation until all of the cells are differentiated.

### Wave velocities

The velocity of a generalized Fisher’s equation of Eq. 15 has the form 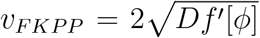. Applied to our system and assuming *∂*_*x*_*ρ* ≈ 0, we obtain the general solution (not assuming *k*_*A*_ = *k*_*B*_)

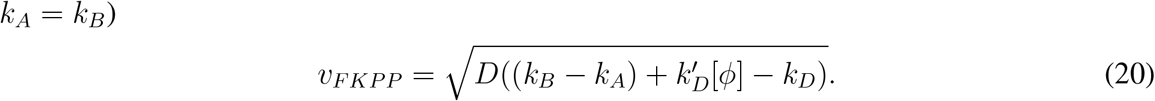

For the specific choice *k*_*D*_ = *α*(*E*[*ϕ*] − *E*_*M*_) (Eq. 18), we obtain

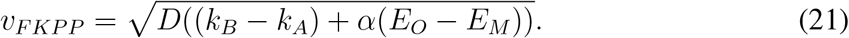

In addition to the Fisher velocity, the cells experience an advection velocity derived from the solution to the force balance equation in Eq. 29. For the cells at the front, this can be written as *v*_*A*_ = *v*(*x* = 0, *t*). The front velocity is then to first approximation given by

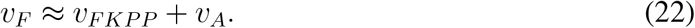

Thus, we now see two main features contributing to the expansion speed: (1) differences in proliferation rate between the cell types and (2) the stiffness gradient across the tissue. First, the FKPP velocity is minimal when the proliferation rates *k*_*A*_ = *k*_*B*_ and increases as the proliferation rate gradient is increases. Secondly, both the advection velocity as well as the FKPP wave front velocity increases with larger differences in stiffness across the system (i.e. upon increasing *E*_*O*_ − *E*_*M*_). In the case of the advection velocity *v*_*A*_, this is due to the mechanical effect of increased pressure gradient across the tissue. In the case of the FKPP velocity *v*_*FKPP*_, this arises from an increase of the differentiation rate with larger stiffness differences as described in Eq. 21.

### Full system

Putting together Eqs. 6, 7, 12 and 13, we obtain the following system of PDEs:

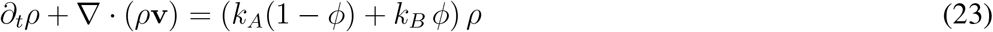

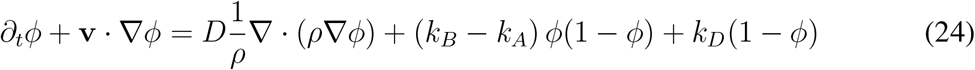

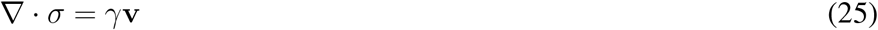

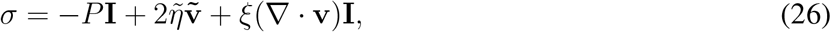

After reduction to 1D and writing *v* = *v*_*x*_, we obtain

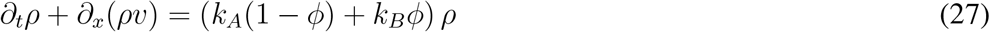

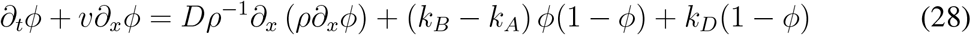

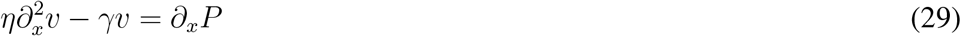

To recapitulate, in the above equation *D* is the diffusion constant, *η* is the viscosity, *γ* is an effective friction term, *k*_*D*_ is the differentiation rate, *P* is the pressure defined in 13, *E* is the stiffness and *ρ*_*h*_ is a homeostatic density. The first equation represents cell number balance due to net proliferation of both cell types as well as advective flows. The second equation describes how the osteoblast fraction *ϕ* changes due to advection, diffusion, cell proliferation, or cell differentiation. Finally, the third equation describes force balance and arises from a balance of viscous stresses with friction and isotropic forces arising from pressure gradients.

We study our system on a bounded spatial domain *x* ∈ [−*L, L*] and take for the initial conditions sigmoidal profiles for *ρ* 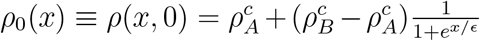and 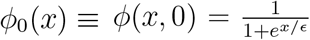 where *ϵ* tunes the sharpness of the profile. As the equation for *v* has no time derivative, we obtain *v*(*x*, 0) as the solution to the non-time dependent Eq. 29 at *t* = 0.

We take Dirichlet boundary conditions for all variables. First, we assume that the cell densities approach their homeostatic values at the boundaries, i.e.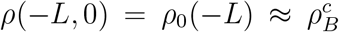 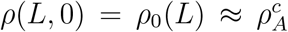. Furthermore, we assume that all cells at the lateral end (where the bone resides) to have differentiated, while none of the cells at the medial end are differentiated, i.e. *ϕ*(−*L*, 0) = *ϕ*_0_(−*L*) ≈ 1 and *ϕ*(*L*, 0) ≡ *ϕ*_0_(*L*) ≈ 0. Finally, we assume that there is no flow of cells at the extreme ends of our system, i.e. *v*(−*L*, 0) = *v*(*L*, 0) = 0. This assumption holds only at short time scales, as the tissue is spatially confined, while on longer time scales the model would need to be extended to take into account the overall growth of the tissue.

We numerically solve this system using a finite difference scheme. Specifically, we use the explicit method for time-stepping and took centered differences for the spatial derivates. We iteratively solve this system in two steps: first we solve the time-independent PDE for *v*(*x, t*) for known profiles of *ρ*(*x, t*) and *ϕ*(*x, t*) at a specific time *t* and then we solve for *ρ*(*x, t* + 1) and *ϕ*(*x, t* + 1) using the explicit scheme.

### Parameter estimates

We estimated the parameters of the model using different methods: inference from experiment data, literature estimates and parameter fitting (2). First, we estimated the carrying capacities of both cell types 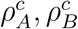from cell density estimates, which we obtained through manual counting of cells in fixed tissue images (Fig. S2D). We estimated the diffusion constant *D* from the cell tracks for each of the three groups of tracked cells by taking the residues obtained from subtracting the average motion of all cells in a group (Fig. S2C). The mean squared displacement of these residues scales roughly linearly with time, which in the case of Brownian motion has a slope of 2*D*. Next, we estimated the stiffnesses *E*_*M*_, *E*_*O*_ from the E14.5 NanoIndenter measurements of the apparent Young’s Modulus (Fig. S2A).

We estimated *τ*, which represents the time scale on which division and apoptosis events lead to homeostasis, to be in the order of 10 hrs. The viscosity *η* and friction coefficient *γ* could also not be directly estimated from our experimental data, but literature estimations are available for both. Moreover, in the inviscid limit (*η* → 0), the cell velocity *v* becomes inversely proportional to *γ* (Eq. 29), such that we can tune this parameter to match the observed cell velocity *v*_*A*_ close to the front. Likewise, we use the measured front velocity *v*_*F*_ to estimate *α*, which from Eq. 22 can be approximated in the following way that depends only on *v*_*F*_ and parameters known from other estimations:

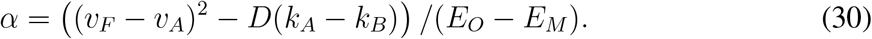

In conclusion, we have two free parameters *γ* and *α* which we fit to the measured velocities *v*_*A*_ and *v*_*F*_ respectively. Thus, the model predictions are in fact the spatial profiles of *ρ, ϕ, V* that can be compared directly to experimental data.

### Analysis of the osteogenic front

#### Definition of the osteogenic front

We start from maximum projections of the live imaging setup for the *Osx1-GFP::Cre* intensity images, and define coordinates *x* running along the medial-lateral axis and *y* in the perpendicular direction. We seek to define an interface that runs vertically along the *y* axis. We set an automated intensity threshold *I* using the Otsu method ((*63*)) and segment the image into connected regions where all pixels have intensity values ≥ *I*. Because of the overall intensity gradient in the *x* direction, there typically is a single largest contiguous region of which the boundary spans the entire vertical axis. Note that in general this boundary could curve in such a way that for some position *y*, there could be multiple points lying on the boundary. Therefore, we first perform an “overhang correction”, by taking the value of *x* that maximizes the size of the contiguous region. The result is then an interface height function *h*(*y, t*) for each frame at time *t*.

#### Quantification of front roughness

There are different ways of defining the roughness of an interface. Here we choose to define roughness through a method commonly used to characterise interfaces in kinetic growth processes (*64, 65*). Specifically, given an interface height function *h*(*y, t*) defined for *y* ∈ {*y*_1_, …, *y*_*n*_}, we first take the (discrete) Fourier transform 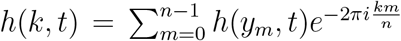.The structure function *S*(*k, t*) is then defined as

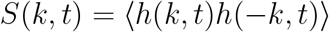

where the average is over replicates of the data (e.g. multiple experiments). Under the scaling hypothesis in interface growth theory, the structure function then scales as (*65*):

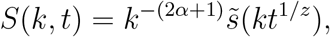

where the dynamic exponent *z* is a constant and

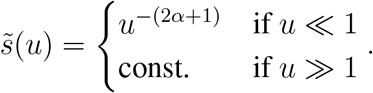

Hence, for sufficiently large times *t, S*(*k, t*) scales as a power law and we obtain the roughness coefficient *α* by fitting our interface function to this power law. Note that in practice, fluctuations at very large *k* (i.e. very small length scales) may deviate from this scaling law. However, these represent fluctuations at spatial scales smaller than that of a single cell. Therefore, in practise we set a cutoff in the wave vector by considering only *k < k*_*max*_ = *L/a* where *L* is the length of the interface and *a* is the typical size of a cell (10 μm).

## Supplementary Figures

### Figure captions

**Supplemental Figure S1.**
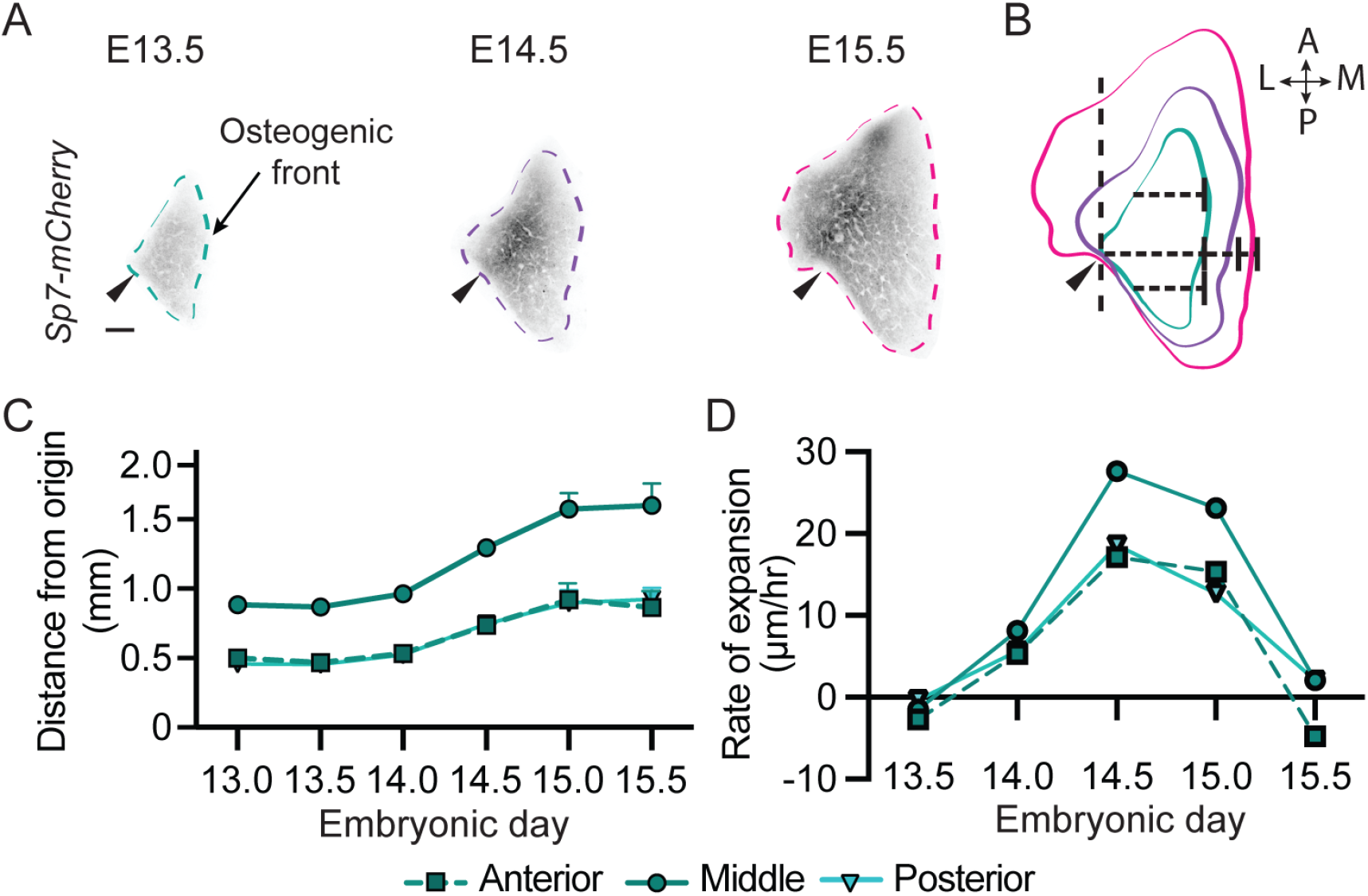
Growth rates of *ex vivo* skull cap explants and *in vivo* skull caps are comparable. **(A)** Frontal bones cropped from flat-mounted skull caps at E13.5, E14.5 and E15.5. Arrows indicate landmarks used to align bones from different samples to one another. **(B)** Example of overlaid bone outlines from different stages. Size measurements were performed at anterior, middle and posterior locations of the bones. A, anterior; P, posterior; L, lateral; M, medial. Scale bar = 500 mm. **(C)** Quantification of medial bone extension between E13.0 and E15.5 at the anterior, middle and posterior positions as indicated in **(C)**. Error bars indicate SD. **(D)** Rates of expansion obtained by comparing subsequent time points for the measurements shown in **(C)**.

**Supplemental Figure S2.**
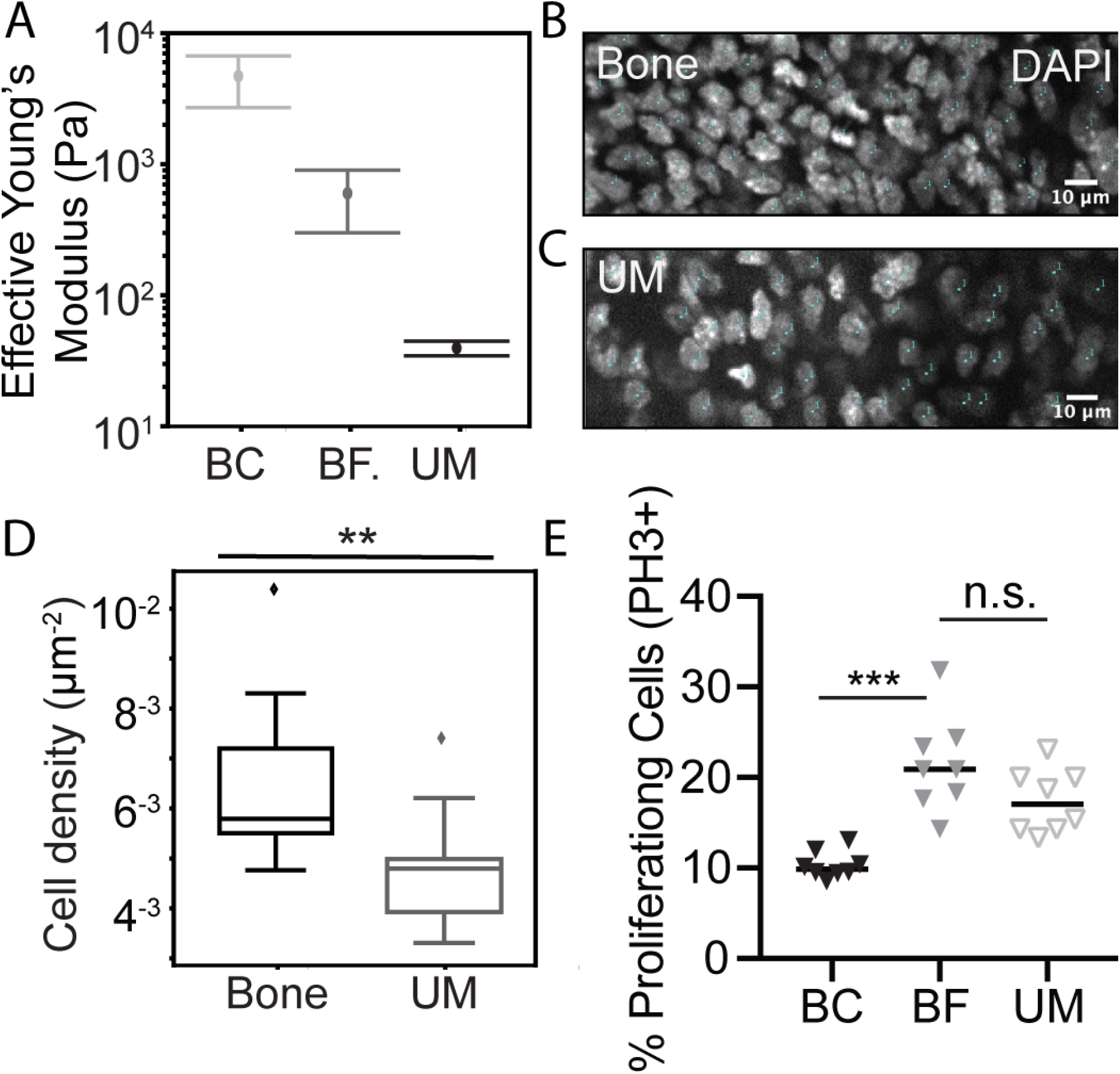
Developing skull bones show regional differences in stiffness, cell density and cell proliferation rates. **(A)** NanoIndenter measurements of the stiffness (apparent Young’s modulus) at the Bone Center (BC), Bone Front (BF) and Undifferentiated Mesenchyme (UM) (indicated in Fig. 1(B)). **(B-C)** Representative DAPI images showing cell nuclei in the bone **(B)** and the undifferentiated mesencyhme **(C). (D)** Quantification of cell densities in the bone (n=22) and the mesenchyme (n=22). Mann-Whitney test, *p <* 0.05. **(E)** Proliferation rates quantified by Ph3+ staining at the Bone Center (N=8, n=360), Bone Front (N=8, n=110) and Undifferentiated Mesenchyme (N=8, n=1177) of the tissue (Mann-Whitney test, *p >* 0.05).

**Supplemental Figure S3.**
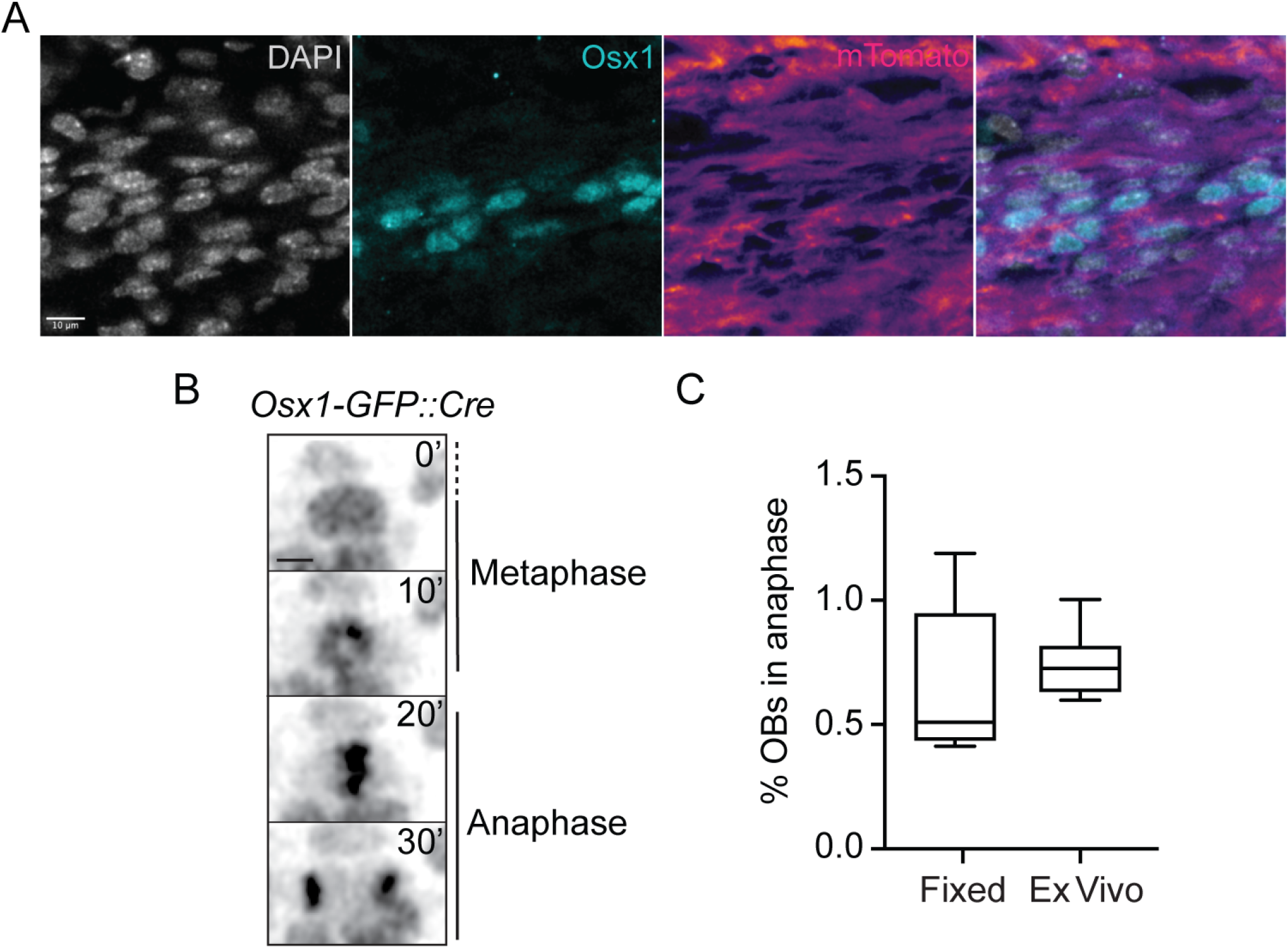
Cell division rates in *ex vivo* experiments match *ex vivo* rates. **(A)** Stainings of cell nucleus (DAPI), osteoblasts (Osx1) and cell membranes (mTomato). **(B)** Stills separated by 10 min of a single *Osx1-GFP::Cre* labelled nucleus during metaphase and anaphase. **(C)** Fraction of osteoblasts in anaphase in fixed tissue images compared to *ex vivo* images.

**Supplemental Figure S4.**
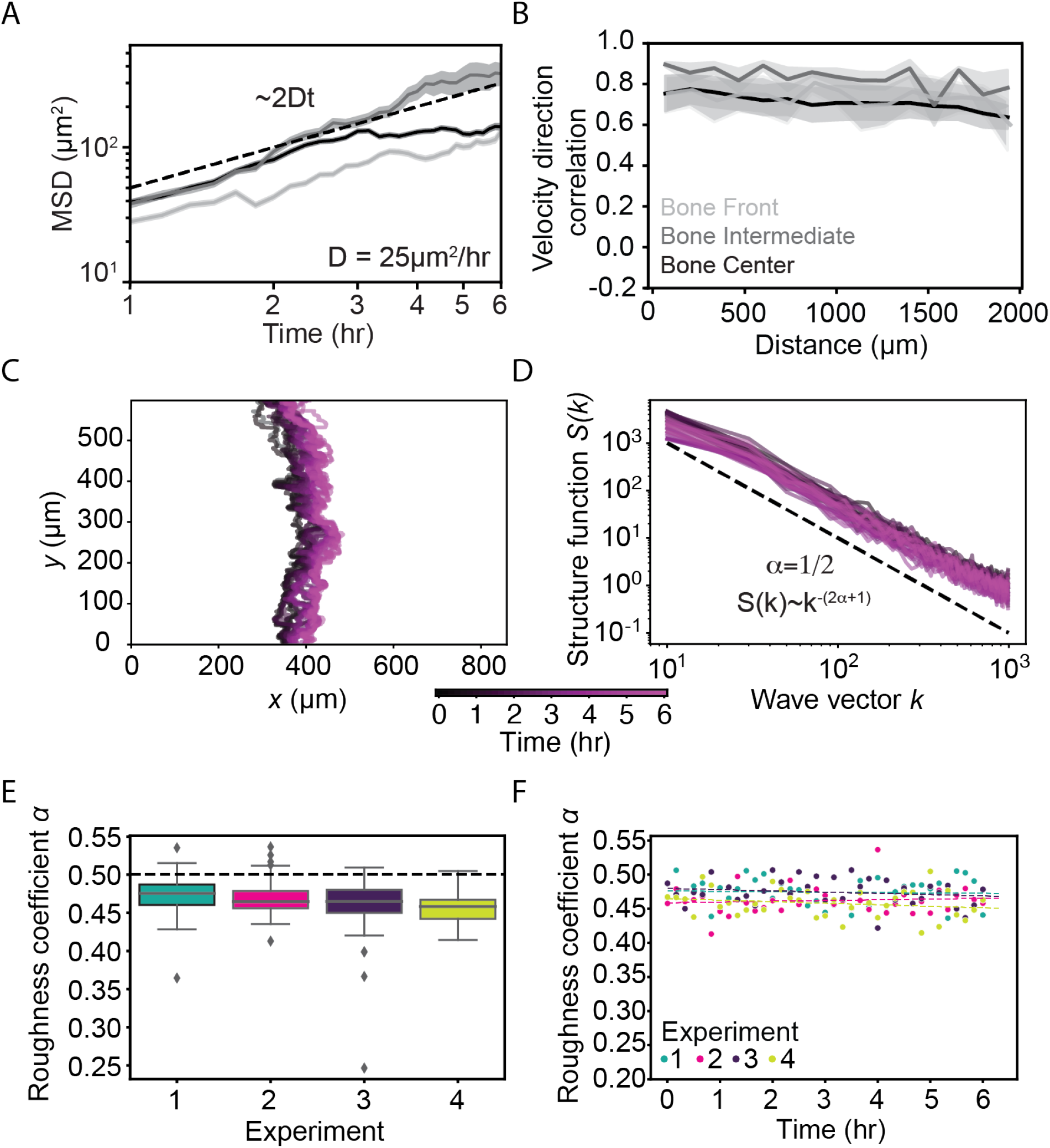
Further quantifications of cell tracks and osteogenic front. **A)** Mean-squared displacements of the relative displacement tracks in (D). Shaded areas show SEM. The dotted line represents diffusive motion with a diffusion constant of *D* = 25 μm^2^*/*h. (B) Normalised velocity correlation at 2 h for four individual movies. Lateral, intermediate and front osteoblast tracks are labelled in light grey, grey and black, respectively. Shading indicated SEM. **(C)** Osteogenic fronts for different time frames of a representative live imaging movie. **(D)** The structure function for the osteogenic fronts shown in **(C)** together scales as a power law, the slope of which defines the roughness coefficient of the interfaces. The dashed line corresponds to a roughness coefficient of 1/2. **(E)** Box plots for roughness coefficients of all frames for four *ex vivo* live imaging experiments. The dotted line marks the value of 1/2. **(F)** Roughness coefficients across time for four *ex vivo* live imaging experiments together with linear fits (dotted lines).

**Supplemental Figure S5.**
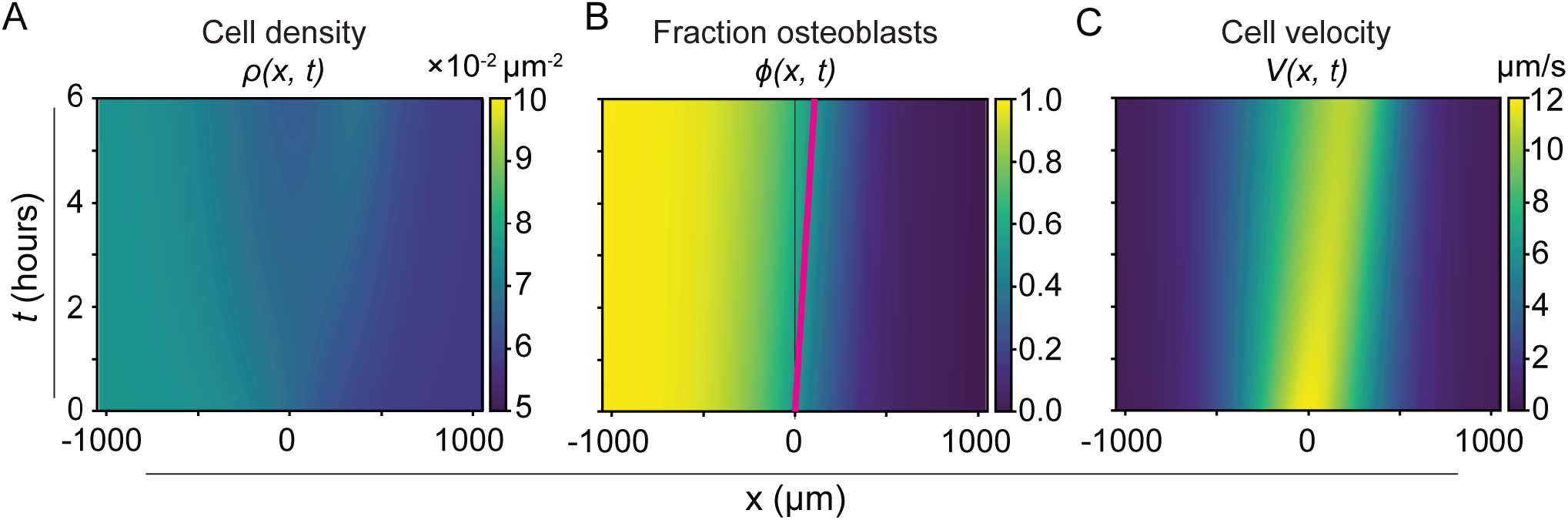
Theoretical model simulations. **(A-C)**Kymographs showing the dynamics of the simulated theoretical model with realistic parameters (see Table 2) for the **(A)** cell density, **(B)** fraction of osteoblasts and **(C)** cell velocity. The magenta line in B indicates a fixed position on the wavefront.

**Supplemental Movie 1. Individual cell borders are complex and dynamic**. Live imaging of mosaic labelling with adenoviral Utr-RFP in wholemount E14.5 skull caps shows complex mesenchymal cell shapes and dynamics.

**Supplemental Movie 2. Bone expansion *ex vivo***. Live imaging of *Osx1-GFP::Cre* labelled skull caps at E13.75.

**Supplemental Movie 3. Oriented division are found at the osteogenic front**. Live imaging of E13.75 *Osx1-GFP::Cre* labelled skull caps.

**Supplemental Movie 4. Few cellular rearrangements at E13.75**. Tracked nuclei show few neighbor exchanges at osteogenic front in live imaged E13.75 *Osx1-GFP::Cre* labelled skull caps.

**Supplemental Movie 5. New differentiation events occur ahead of the osteogenic front**. Live imaging of E13.75 *Osx1-GFP::Cre; R26RmTmG* reporter mice showing nuclear only label ahead of an osteogenic front labelled with both membrane and nuclear GFP.

